# Polarized ATP synthase in synaptic mitochondria induced by learning and plasticity signals

**DOI:** 10.1101/2024.11.26.625359

**Authors:** Yi Hu, Xi Wang, Kaiyuan Liu, Songhai Xiong, Ji-Song Guan, Hong Xie, Min Gu

## Abstract

MINFLUX represents a state-of-the-art single-molecule localization microscopy technology that surpasses the conventional diffraction limit, enabling the visualization and analysis of nanostructures with exceptional precision. Despite its potential, the utilization of MINFLUX has been primarily confined to in vitro cultured cell environments. In this research, we have refined the sample preparation protocols to facilitate 3D MINFLUX imaging in fixed brain tissue sections, focusing on the mitochondrial distribution within dendritic spines and engram cells of the dentate gyrus. Probing single molecules *in vivo* reveals an interesting finding: the reorganization of mitochondrial inner membrane proteins during synaptic plasticity within dendritic spines of engram cells. Utilizing 3D MINFLUX nanoscopy, we identified a significant redistribution of α-F1-ATP synthase, which correlates with learning related activities. This redistribution implies a pivotal function for polarized ATP synthesis in the vicinity of the postsynaptic zone, which may modulate both short-term and long-term synaptic modifications, thereby influencing synaptic plasticity and memory consolidation. Furthermore, dual-color 3D MINFLUX imaging has uncovered distinct patterns of mitochondrial reorganization involving both the inner and outer membranes within dendritic spines. These patterns, induced by plasticity signals, persist for up to 12 hours in neuronal cultures. This distinction highlights the presence of distinct regulatory mechanisms governing mitochondrial proteins during synaptic plasticity. These results offer new insights into the molecular mechanisms underlying synaptic plasticity and underscore the transformative potential of 3D MINFLUX imaging for studying neuronal processes in the brain.

## Introduction

Memory engram cells are crucial for the formation and storage of long-term memories, serving as the cellular substrate underlying the brain’s cognitive functions^1–6^. These specific neuronal populations undergo significant alterations during learning, forming tightly interconnected networks that mediate memory consolidation and retrieval^7–9^. Synaptic plasticity is fundamental to the stability of memory engram networks. It enhances the connectivity between engram cells by modifying the structural and electrophysiological properties of synapses^10–12^. These adaptations are crucial for stabilizing memory traces and ensuring effective reactivation of the memory network upon exposure to reminder cues^13^.

Mitochondria play a critical role in supporting synaptic plasticity by providing the energy necessary for synaptic transmission and neuronal function^14^. As the primary site of ATP production, mitochondria ensure a consistent supply, especially during periods of high energy demand such as synaptic transmission and plasticity^15^. The energy consumption at the synapse is substantial, and mitochondria, strategically located near synapses^16–18^, are essential for maintaining synaptic function by rapidly supplying ATP. Despite their crucial role, current research on mitochondria has primarily focused on overall mitochondrial function rather than detailed protein-level analysis within specific cellular contexts.

Recent advancements in single molecue localization microscopy(SMLM) techniques, such as MINFLUX, offer high-resolution insights into cellular structures and molecular dynamics^19,20^. MINFLUX excels in providing nanometer-scale resolution and microsecond-range tracking, making it particularly valuable for studying mitochondrial membrane proteins^21^. However, most studies using MINFLUX have been conducted in controlled environments like cultured cells, leaving a knowledge gap regarding mitochondrial protein distribution in native biological tissues.

This study aims to address this gap by utilizing 3D MINFLUX to investigate the spatial distribution of mitochondrial proteins within engram cell synapses. Our research discovered a redistribution of the mitochondrial inner membrane protein ATP5a in dendritic spines of memory engram cells, concentrating in the direction of synaptic contact sites. Additionally, through in vitro cell culture and chemical LTP induction to simulate the learning process, we found that this redistribution phenomenon of synaptic mitochondrial inner membrane α-F1-ATP synthase (ATP5a) occurs in both spine synapses and shaft synapses. In contrast, the outer membrane protein: translocase of outer mitochondrial membrane 20 (TOMM20) did not exhibit such a polarized redistribution. These findings suggest that the spatial distribution of mitochondrial inner membrane proteins may play an important role during the learning process. Optimizing their spatial distribution may enhance energy-consuming biological processes associated with synaptic plasticity. This result also provides a nanoscale explanation for mitochondrial involvement in learning and memory-related processes, enriching our understanding of the mechanisms underlying brain cognitive functions.

## Results

### Nanoscale distribution of mitochondrial ATP5a in dendrites of memory engram cells revealed by MINFLUX nanoscopy

To investigate the function of mitochondria in the synaptic plasticity of memory engram cells, activity-dependent labeling was performed in the hippocampal region of cfos-CreER mice. The TRAP (Targeted Recombination in Active Populations) system^22,23^, a widely used tool in neurobiology, was employed for activity-dependent labeling. AAV vectors carrying mCherry flanked by loxP sequences were injected into the hippocampal region of the mice. To label both the engaged memory engram cells and the nearby non-engaged neuronal populations, a 1:1 mixture of the virus carrying a fluorescent EGFP sequence driven by the CamKII promoter was used, allowing for activity-dependent marking while simultaneously labeling nearby non-engaged neurons(Fig. 1a). Following viral infection and during learning-induced neuronal excitation, the expression of the cfos gene drove the transcription of CreER. Tamoxifen, administered 24 hours prior, bound to CreER, removing its spatial hindrance and allowing its nuclear entry. This facilitated recombination at loxP sites, leading to the expression of mCherry and marking the memory engram cells (Fig. 1b). A timeline of the experimental procedure is illustrated in Fig. 1c. After a two-week recovery period post-injection, the mice underwent intraperitoneal injection of Tamoxifen(150 mg/kg), followed by contextual fear conditioning training 24 hours later. During the training process, activated neurons were marked. Three days post-training, The mice were sacrificed, and brain tissue was collected and processed for sectioning and subsequent analysis. Confocal imaging revealed the presence of mCherry-labeled memory engram cells in the dentate gyrus, while EGFP marked the surrounding non-engaged neurons(Fig. 1d;Supplementary Fig.1a,b). Consistent with previous findings^7^, structural analysis revealed that memory engram cells exhibited significantly more dendritic spines compared to non-engaged cells(Fig. 1e). Additionally, dendritic spine width was statistically greater in the memory engram cell populations, indicating significant structural plasticity and enhanced synaptic connectivity(Fig. 1f).

**Fig 1.**
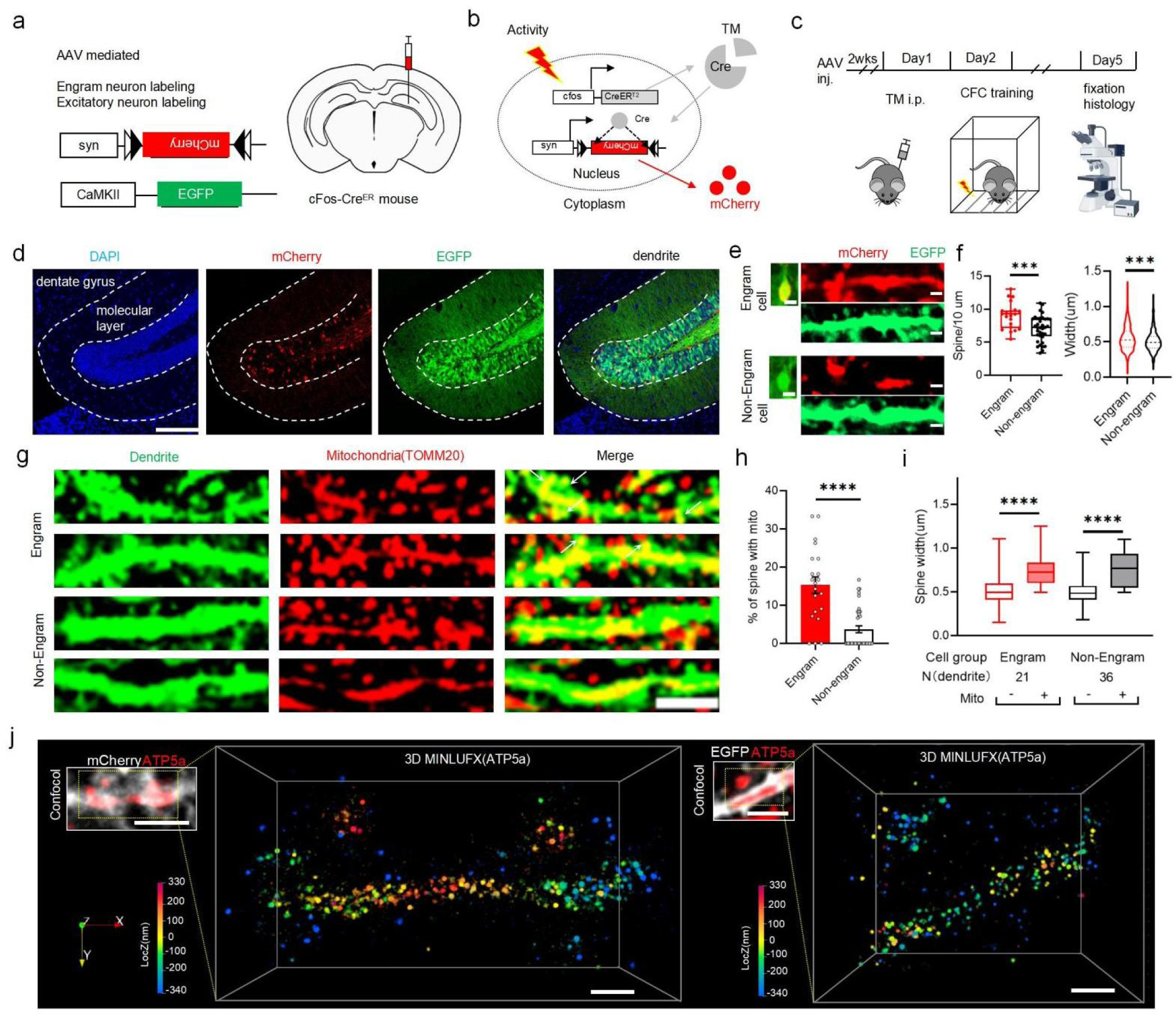
3D MINFLUX Imaging of Mitochondrial Protein Distribution in Memory Engram Cells in Brain Tissue Sections. **a,c** Labeling of memory engram and non-engram cells in the hippocampal DG region by AAV viral vectors **a** Expression of EGFP protein driven by the CamKII promoter for labeling excitatory neurons in the hippocampal DG region. mCherry protein expression driven by the Syn promoter, with loxp sequences, labels activity-dependent memory engram cells. Both viruses were mixed at a 1:1 ratio and injected into the hippocampal DG region of cfos-creER mice.**b** Overview of the TRAP technique. By combining TRAP with tamoxifen (TM) injection, active neurons within a specific time window can be fluorescently labeled.**c** Schematic representation of the behavioral protocol for labeling in mice. **d** Representative confocal images showing memory engram and non-engram cells. The blue channel represents DAPI. The red channel shows mCherry-labeled memory engram cells, while the green channel highlights EGFP-labeled excitatory neurons. Scale bar = 200 μm. **e,f** Observation and statistical analysis of dendritic spine density in memory engram and non-engram cells. Cells with co-localized red and green signals, along with dendrites, are classified as memory engram cells, while cells with only green fluorescence are categorized as non-engram cells (n-engram cells = 21, n-non-engram cells = 36), Two-tailed unpaired t-test, p=0.007 for spine density quantification and p= 0.004 for spine width quantification. **g,i** Observation and statistical analysis of mitochondrial distribution in dendrites and dendritic spines. Mitochondria were labeled by immunofluorescent staining of TOMM20 which was first labeled using primary antibodies conjugation with AF647. White arrows indicate mitochondria located within dendritic spines. Scale bar = 5 μm. **h** Probability statistics of mitochondrial localization within dendritic spines of memory engram and non-engram cells, Two-tailed unpaired t-test, p<0.0001.**i** Morphological differences of dendritic spines containing mitochondria between the two cell groups, compared with dendritic spines devoid of mitochondria, Two-tailed unpaired t-test, p(Engram cell mito+ vs Engram cell mito-)<0.0001, p(non-Engram cell mito+ vs non-Engram cell mito-)<0.0001, p(Engram cell mito+ vs Engram cell mito-)<0.0001. **j** 3D MINFLUX imaging of ATP5a protein in mitochondria located in mCherry-positive dendrite(left) and EGFP positive dendrites(right). The z-axis is presented in a color-coded manner. ATP5a was first labeled using a primary antibody, followed by secondary antibody conjugation with AF647. Scale bar for confocal images = 2 μm, scale bar for 3D MINFLUX images = 500 nm. Error bar shows S.E.M. *P < 0.05, ***P < 0.001, ****P < 0.0001.

Having successfully targeted the memory engram cell populations, we next examined the distribution of mitochondria and its relationship with the maturation of dendritic spines in these cells. Mitochondria were labeled with an anti-TOMM20 antibody conjugated to the AF647 dye, and their distribution within dendrites and dendritic spines was observed via confocal microscopy(Fig.1g;Supplementary Fig.2a,b). The results showed that the percentage of mitochondria presence in dendritic spine is bigger in memory engram cells populations, with mitochondria being preferentially located in larger spines, which are typically associated with more mature synaptic plasticity processes(Fig. 1h-i). Next, we explored the potential significance of the mitochondria’s distribution in dendritic spines. Mitochondria contain numerous membrane proteins on both the inner and outer membranes, which intricately mediate their functions^24,25^. However, traditional confocal microscopy, due to its resolution limitations, is unable to resolve the distribution of these proteins. To capture the finer details of mitochondrial changes within synapses, we turned to 3D MINFLUX imaging. Before conducting MINFLUX imaging on brain sections, several considerations were made to optimize tissue preparation. Due to the thickness of brain tissue compared to cultured cells, sectioning parameters were adjusted by increasing the oscillation frequency of the microtome and reducing the blade advancement speed, which allowed for the production of thinner sections of brain tissue (10-15 µm). Additionally, to reduce non-specific antibody binding, which could lead to excessive fluorescence signal, two key adjustments were made: the blocking time with BSA was extended to minimize non-specific binding; and during the immunohistochemistry process, the PBS wash time and volume were optimized to further reduce background fluorescence.(Supplementary Fig.3a-b). Following traditional mounting protocols of MINFLUX imaging in cultured cells^21^, GLOX buffer that contains 30mM MEA was used for later imaging studies.

As shown in Fig. 1j, 3D MINFLUX imaging revealed the distribution of ATP5a proteins (was labeled with primary antibody, followed by secondary antibody conjugated to AF647) within the dendrtite and the spines of the marked memory engram cells and non-engram cells(Supplementary Fig. 3c). Additionally, we also achieved 2D and 3D dual-color MINFLUX imaging in brain sections, where TOMM20 was labeled with primary antibody conjugated to AF647, and ATP5a was labeled with primary antibody followed by a secondary antibody conjugated to FL680 dye (Supplementary Fig. 3d). Using DBSCAN clustering and spatial analysis^26^, the distribution pattern of ATP5a molecules was analyzed(Supplementary Fig.4a-b). The spatial resolution of protein localization was confirmed with XYZ axis precision of 6-7 nm (Supplementary Fig.4c). The nearest neighbor distance distribution of ATP5a molecules within memory engram cells revealed several distinct peaks, with a major peak 21.27 nm, and additional peaks at 52.25 nm, 72.16 nm, 87.65 nm, and 116.4 nm(Supplementary Fig.4d), validating the high spatial resolution of MINFLUX and confirming its applicability for observing molecular organization at the nanoscale.

### Polarized redistribution of mitochondrial inner membrane protein ATP5a after learning-induced synaptic plasticity

To explore the role of mitochondrial reorganization during learning, particularly within synapse-associated dendritic spines, the distribution of ATP5a in spine areas with pre-synaptic contact was specifically examined. Confocal microscopy revealed a tight spatial relation between the localization of mitochondria and pre-synaptic markers synaptophysin(syn), suggesting that mitochondria play a significant role in synaptic function(Fig. 2a,2c). Further Analysis of 3D MINFLUX data indicated that ATP5a exhibited preferential clustering towards the direction that were enriched with syn signals, while this redistribution was absent in the control group (EGFP-labeled non-engram cells)(Fig. 2b,2d). To quantify this phenomenon, DBSCAN clustering analysis was performed to locate the labeled molecules and the local density of each ATP5a molecules were then calculated(Fig. 2e), revealing a higher local density of ATP5a molecules in syn positive directions. This was further confirmed by radial measurements of local density using segmentation approach(Fig. 2f), which indicated a marked increase in ATP5a local density in syn direction(syn+) in dendritic spine regions memory engram cells compared to non-engram cells(Fig. 2g). These results suggest that the molecular reorganization of ATP5a within dendritic spines is closely tied to synaptic plasticity during the learning process(Fig. 2h).

**Fig 2.**
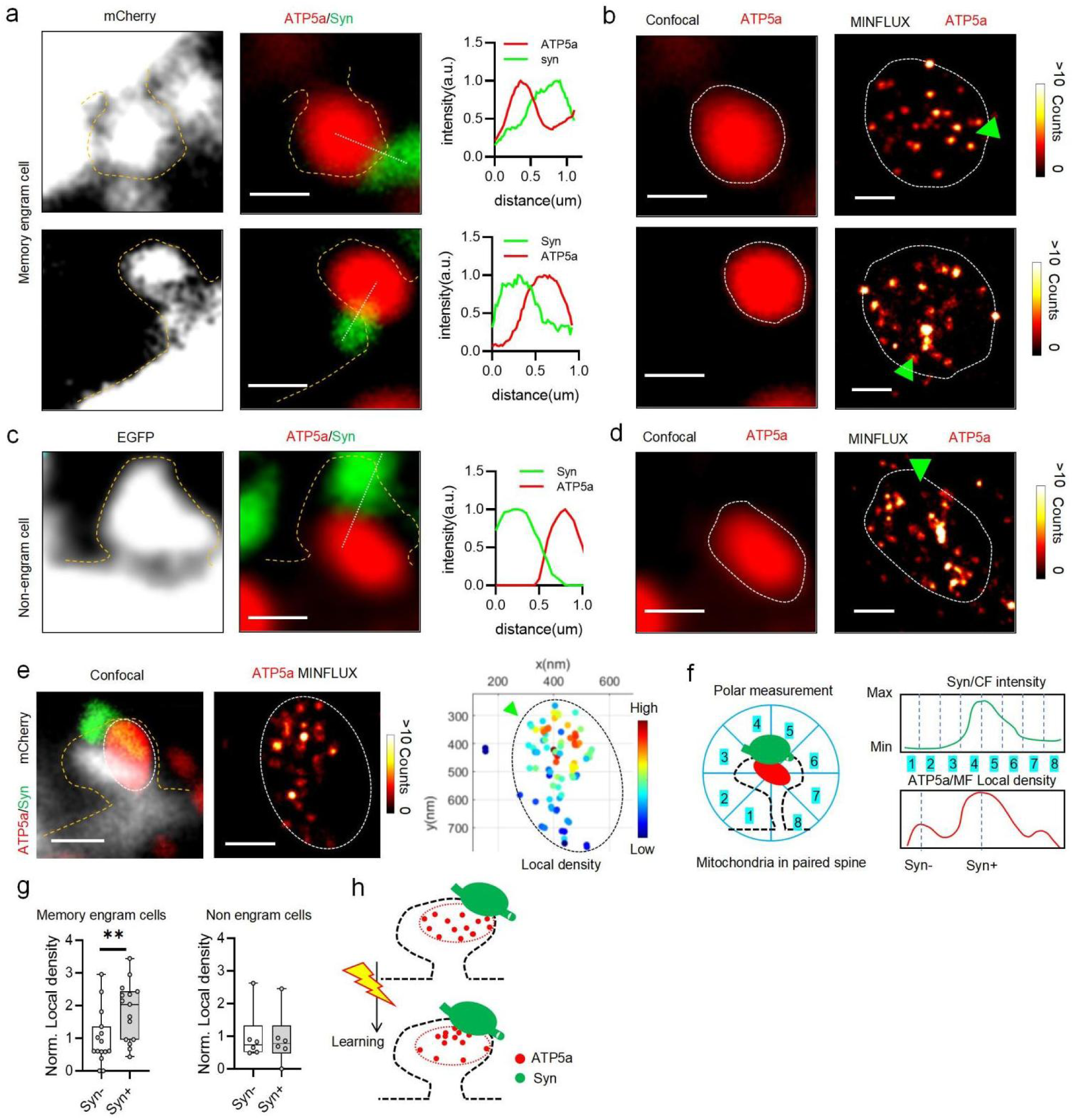
Polarized Redistribution of mitochondria inner membrane protein ATP5a in dendritic spine of DG memory engram cell. **a,d** Super-resolution mapping of ATP5a distribution, a mitochondrial inner membrane protein, in dendritic spines of memory engram cells and non-engram cells in the molecular layer.**a,b** Representative MINFLUX imaging of ATP5a in dendritic spines of memory engram cells. Left: mCherry-labeled dendritic spines (gray channel) outline the structural morphology. Middle: dual-channel images showing ATP5a (red) and synaptophysin (green) staining, with intensity profiles indicating tight colocalization. Right: Z-projection of 3D MINFLUX data for mitochondrial ATP5a in the region indicated, where green arrows highlight regions of high syn intensity correlating with enriched MINFLUX localizations. Confocal scale bar, 400 nm, MINFLUX scale bar, 200nm. **c, d** Representative MINFLUX imaging of ATP5a in dendritic spines of excitatory neurons labeled via CamKII promoter-driven EGFP expression. Confocal scale bar, 400 nm, MINFLUX scale bar, 200nm. **e, f** Analysis pipeline for correlating ATP5a MINFLUX signal intensity with syn intensity in dendritic spines. **e** Workflow begins with selecting regions of interest (ROIs) containing mitochondria within dendritic spines of an engram cell for MINFLUX data acquisition. Clusters of ATP5a molecules were identified using DBSCAN-based clustering, and local densities were calculated to assess ATP5a molecule distributions. Scale bar, 200 nm. **f** Local density and synaptic intensity data were measured using an 8-segment division centered on the mitochondrial staining centroid. Regions with the highest synaptic intensity (syn+) and the lowest synaptic intensity (syn−) were compared to analyze differences in ATP5a molecular density. **g** Spatial redistribution of mitochondrial ATP5a relative to synaptic regions in dendritic spines of memory engram cells versus non-engram cells. Statistical analysis revealed significant differences in the spatial distribution of ATP5a between these two groups. (n-memory engram cell = 15 from 6 mice; n-non-engram cell = 6 from 3 mice; two-tailed paired t-test, p = 0.0018). **h** Proposed molecular model of ATP5a reorganization in dendritic spines of memory engram cells during learning-related processes. This model summarizes observed spatial changes in mitochondrial inner membrane protein distribution. Error bars represent S.E.M.**P < 0.01.

### Chemical-induced long-term potentiation (cLTP) triggers ATP5a reorganization in dendritic spines

To further explore the relationship between synaptic plasticity and mitochondrial inner membrane protein reorganization, we conducted MINFLUX imaging experiments in cultured cortical primary neurons. AAV mediated expression of membrane GFP was conducted in order to visualaize the morphology of he neuron(Fig.3a). To simulate learning-related processes in vitro, chemical-induced long-term potentiation (cLTP) was used^17^(Supplementary Fig.5a-b). DIV17-21 neurons were exposed to a short 5min duration of 100 μM glycine stimulation, after which they were fixed and immuno-stained for ATP5a and syn. Confocal microscopy revealed that mitochondrion in dendritic spines were located near pre-synaptic contact sites in both spine synpases and shaft synapses(Fig.3b). Consistent with *in vivo* findings, 3D MINFLUX imaging revealed that ATP5a molecules in dendritic spines were significantly redistributed following cLTP induction. ATP5a clustered near synaptic contact sites in cLTP-stimulated neurons, whereas this redistribution was not observed in the control group(Fig. 3c-f). ATP5a MINFLUX localizations were collected in the shaft spine region and its adjacent areas with weaker pre-synaptic signals(Fig. 3g). The local density of each identified ATP5a molecule was calculated to quantify its distribution(Fig. 3h). Further analysis of local density maps ATP5a of shaft spine region indicated a strong correlation between ATP5a local density and pre-synaptic signal strength, particularly in syn high signal regions of cLTP group(Fig. 3i-k). Additionally, the temporal dynamics of ATP5a redistribution were assessed through a time-course experiment. Neurons were fixed and analyzed at 30 minutes, 12 hours, and 3 days post-cLTP induction(Fig. 3l). The results showed that ATP5a reorganization persisted for at least 12 hours after cLTP induction, although the effect diminished by 3 days(Fig. 3m), suggesting a transient but significant role for ATP5a in synaptic plasticity and the difference between neural networks *in vitro* and memory engram networks *in vivo*.

**Fig 3.**
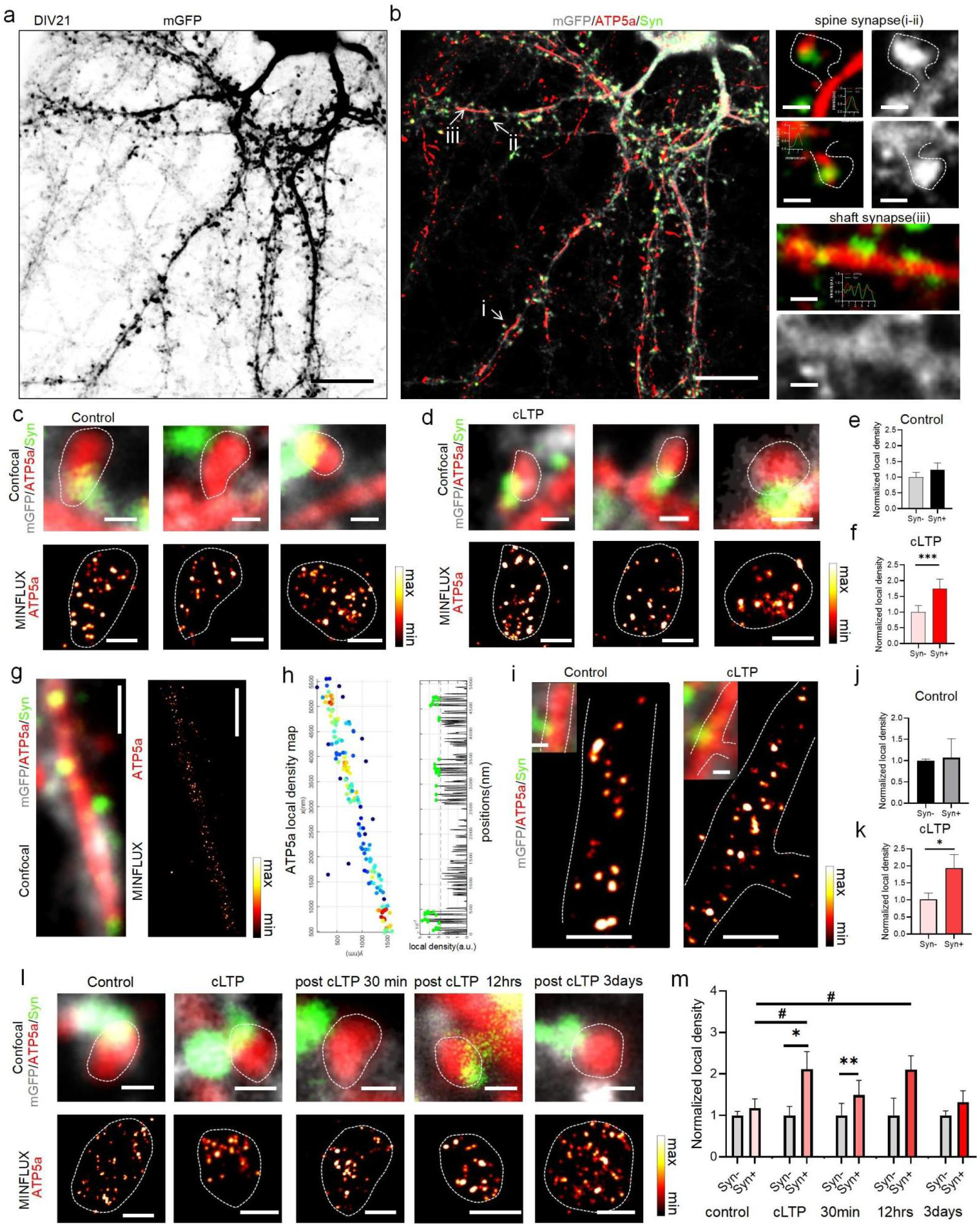
Polarized redistribution of mitochondria inner membrane protein ATP5a during cLTP process. **a.** Representative images of DIV21 cultured primary cortical neurons expressing mGFP. Scale bar: 50 μm. **b.** Left:Mitochondria were labeled using primary antibodies against ATP followed by secondary antibodies conjugated with AF647. Synapses were labeled using primary antibodies followed by secondary antibodies conjugated with AF555, right: Representative images showing postsynaptic mitochondria in spine synapses and shaft synapses. Inserted figures are line profiles of ATP5a (red) and Syn (green) confocal signals across the postsynaptic sites in spine and shaft synapses. **c, f** Reorganization of mitochondrial ATP5a in dendritic spines under cLTP and control conditions. **a** Confocal images show mitochondrial ATP5a (red, AF647) and synaptophysin (green, AF555) in dendritic spines of cultured neurons. The white dashline indicates the mitochondria area. The bottom panel shows z-projection of MINFLUX data for ATP5a localization. Confocal scale bar: 400 nm; MINFLUX scale bar: 200 nm. **e, f** Quantification of ATP5a molecular local density enrichment in synaptic regions in control and cLTP conditions. Data were statistically analyzed using a two-tailed paired t-test, p = 0.0005. (n-cLTP = 25 neurons, n-ctrl = 24 neurons from 3 separate neuronal cultures). **g, k** Reorganization of mitochondrial ATP5a in shaft and spine regions. **g** representation of z projection of 3D MINFLUX imaging of ATP5a in the shaft spine regions. **h** Distribution of ATP5a molecule local density along the y-axis. The black dashed line indicates the mean molecular density plus one standard deviation (SD). Regions above this threshold are marked with green spheres. **i** 3D MINFLUX imaging of ATP5a in the shaft spine regions in control group and cLTP group. j**,k**Quantification of ATP5a molecular local density enrichment in synaptic regions in control and cLTP conditions. Data were statistically analyzed using an unpaired t-test, p = 0.0257. (n-cLTP = 10 neurons, n-ctrl = 5 neurons from 3 separate neuronal cultures). **l,m** Distribution of mitochondrial ATP5a protein in dendritic spines at different time points after cLTP induction. **l** Top: Immunohistochemistry results showing mitochondrial ATP5a (red) and adjacent synaptophysin (syn, green) in dendritic spines. Bottom: Z-axis projection of MINFLUX data showing ATP5a localization in the same dendritic spines. Confocal scale bar: 400 nm; MINFLUX scale bar: 20 nm. **m** Quantification of local density enrichment of ATP5a in synaptic regions. Data are presented for cLTP group syn+ versus syn− (p = 0.0185), post-cLTP 30 min group syn+ versus syn− (p = 0.0037), cLTP syn+ versus control syn+ (p = 0.0399), and post-cLTP 12 hr syn+ versus control syn+ (p = 0.0249). */^#^P < 0.05, **P < 0.01. Error bars represent S.E.M.

### Lack of cLTP-induced reorganization of the outer membrane protein TOMM20

Mitochondria consist of both inner and outer membrane proteins with distinct functional roles. While the reorganization of the inner membrane protein ATP5a was observed during synaptic plasticity, it remained unclear whether outer membrane proteins such as TOMM20 exhibit similar changes. To address this, we performed dual-labeling of mitochondrial inner and outer membrane proteins, with ATP5a and TOMM20 immuno-labeled with different fluorophores (FL640 and FL680, respectively)(Supplementary Fig.6a-d),The localization precision of both proteins was quantified at approximately 6-7 nm(Supplementary Fig.6e), validating the high spatial accuracy of the MINFLUX system. 3D MINFLUX imaging revealed that, unlike ATP5a, TOMM20 did not exhibit any specific redistribution in response to cLTP induction. (Fig. 4a-d;Supplementary Fig.7a-c). These results suggest that TOMM20, as an outer membrane protein, does not undergo the same dynamic reorganization as ATP5a during synaptic plasticity, highlighting the differential roles of mitochondrial membrane proteins in synaptic remodeling.

**Fig 4.**
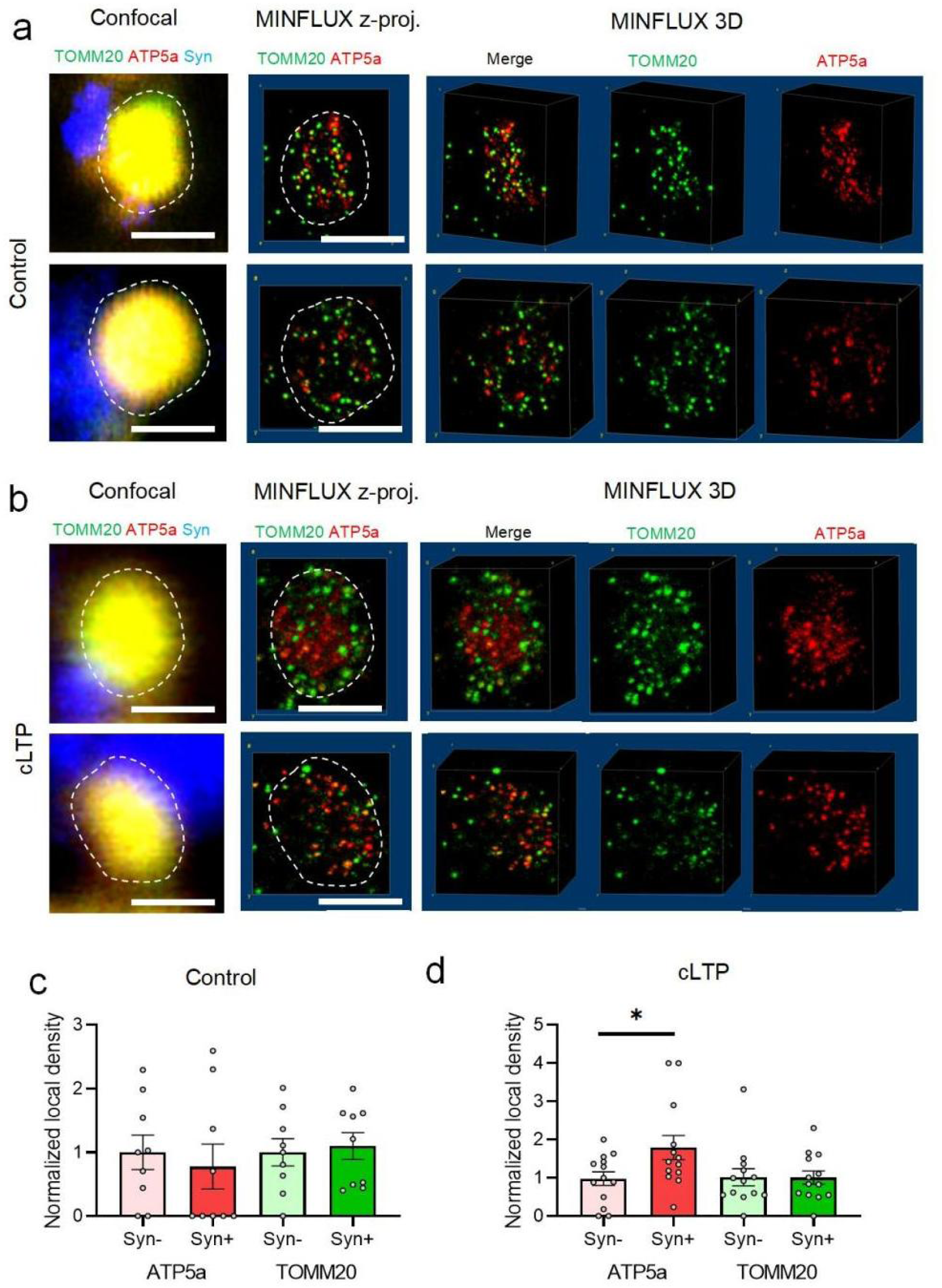
Distinct redistribution patterns of inner and outer mitochondrial membrane proteins in dendritic spines. **a,b** 3D dual-color MINFLUX imaging results of dendritic spines.**a** Example from the control group. Confocal images show ATP5a protein (red channel), TOMM20 protein (green channel), and synaptic marker (blue channel) from virus-induced syn expression. Right panel: 3D MINFLUX data reconstructed using ImageJ, with left showing the z-axis projection and right showing the 3D rendering. Pseudo-coloring follows the same scheme as in confocal images. **b** Example from the cLTP group, with the same imaging format as in (e). Scale bar: 400 nm. **c,d** Comparison of local density differences of ATP5a and TOMM20 proteins between syn+ and syn-regions in the control and cLTP groups. Red bars represent ATP5a data, while green bars represent TOMM20 data. Local density calculations for dual-color imaging are provided in the supplementary materials. n-cLTP = 20 neurons, n-control = 9 neurons. Two-tailed paired t-test, p = 0.0412. *P < 0.05. Error bars represent standard error of the mean (S.E.M.).

## Discussion

Recent studies have elucidated the cellular mechanisms underlying the formation and maintenance of memory engrams in the brain, primarily using immediate early gene-based labeling systems combined with in vivo imaging, optogenetics, and chemogenetics^9,27,28^. Through observation of memory engram cells in the hippocampal DG region, researchers have found that the dendritic spine density of memory engram cells is significantly higher than that of non-engram cells^29^. Furthermore, by expressing light-sensitive channel proteins in the entorhinal cortex (EC) and using light to activate the projections to the DG, researchers observed that memory engram cells in the DG produced higher excitatory postsynaptic currents^29^. These findings indicate a specific increase in synaptic strength in the memory engram cell population. The plasticity of synapses after learning is dependent on new protein synthesis and this process at the synaptic structures of neurons is highly energy-demanding, which highlights the critical role of mitochondrial function in synaptic plasticity. in line with this, studies have found that synaptic activities promote the presence of mitochondria in the dendritic protrusions^16^ and knockout of PINK1, a regulator of mitochondria dynamics, shows reduction of mitochondria localizations in dendritic spine and creates defect in learning and memory. In our study, we utilized the 3D MINFLUX nanoscopy technique, which offers exceptional single-molecule localization capabilities, to observe that the ATP5a protein on the inner mitochondrial membrane in the dendritic spines of memory engram cells in the mouse hippocampus undergoes redistribution. This suggests that, after learning, mitochondria actively participate in the plasticity changes occurring at the dendritic spines.The polarized distribution of ATP5a may enable its rapid diffusion to synaptic sites, promoting synaptic plasticity, including dynamic changes in actin, AMPA receptor trafficking, and local protein translation processes^14^. The redistribution phenomenon observed in the shaft spines further supports this conclusion. It is known that in excitatory principal neurons, a certain proportion of glutamatergic synapses remain located in the dendritic shaft^30^. The maturation process of shaft spines, which protrude to form more mature spines after learning, depends on F-actin remodeling^31^, and optimized ATP5a protein distribution may further promote this process.

Interestingly, in the *in vitro* cLTP model used to simulate learning, we observed that the ATP5a redistribution phenomenon only lasted for up to 12 hours. However, in memory engram cells, the redistribution was observed the third day after learning. This discrepancy may be due to several factors. First, after memory formation, the brain undergoes consolidation processes such as hippocampus-mediated replay^32,33^, which could require a longer duration of the polarized redistribution process to further support energy needs. Another possibility is that cLTP induces more mitochondria to enter the spines, but these later-entering mitochondria may not receive the most direct stimuli and thus fail to undergo redistribution. At the same time, we also observed that the outer mitochondrial membrane protein TOMM20 did not undergo specific redistribution during the synaptic plasticity process. This suggests that the distribution of inner and outer mitochondrial membrane proteins is regulated by different mechanisms. TOMM20 is a crucial component of the mitochondrial outer membrane translocase complex, responsible for importing mitochondrial proteins from the cytoplasm into the mitochondrial outer membrane. Therefore, this process may not be regulated by synaptic plasticity.

MINFLUX, a localization concept proposed in 2017^34^, has pushed 3D multicolor single molecule imaging and tracking to unprecedented levels, achieving single-digit nanometer precision and approximately 100 µs time resolution, with a scalable field of view^19^. In this study, we pioneered the use of MINFLUX 3D imaging in biological tissues, particularly in neuroscience research. *In vitro* cultured neuron systems often differ greatly in network characteristics from those in the biological brain, making *in vivo* tissue observations essential for obtaining more accurate biological insights. Additionally, synaptic plasticity occurs at the nanoscale level of synaptic connections. Previous studies using STED/STORM super resolution imaging research have provided valuable insights such as the aligned nanomodules of pre-and postsynaptic proteins that contributes to structural plasticity^35^ and the existence of trans-synaptic molecular nanocolumns^36^. Given these findings, we believe MINFLUX can further enhance our understanding of nanoscale biological processes involved in learning, as well as the abnormalities in synaptic regulation seen in neurological diseases, thus providing deeper insights into disease mechanisms.

## Materials and methods

### Mouse subjects

In this study, cFos-CreER mice (stock number 021882, The Jackson Laboratory) and wild type C57BL/6 mice. All animals were housed in the Laboratory Animal Facility at ShanghaiTech University, which is accredited by the Association for Assessment and Accreditation of Laboratory Animal Care International. under a 12-hour light/12-hour dark cycle (9 a.m.–9 p.m.) with unrestricted access to food and water. The experimental mice were males, aged 3–4 months. All mice were allowed to recover at least one week from virus injection before any behavioral tasks. All animal protocols and experiments were reviewed and approved by ShanghaiTech University (license number 20201218002), following the Guide for the Care and Use of Laboratory Animals and in accordance with Chinese law (Laboratory Animal-Guideline for Ethical Review of Animal Welfare, GB/T 35892). This study adheres to all relevant ethical regulations for animal testing and research and received ethical approval from the Scientific Research Ethics Committee at ShanghaiTech University.

### Mouse Virus Injection

Mice were first anesthetized with 1.5% isoflurane. The skin on the top of the head was disinfected and then incised along the midline. A craniotomy was made in the skull and the virus was injected at the following coordinates: AP-2.0mm, ML ±1.5mm, DV-2mm. 100 nl of AAV2/9-DIO-mCherry and AAV-CamkII-EGFP was injected at a rate of 50 nl/min. The needle was left in place for 2 minutes following the injection before being withdrawn. The skin along the midline of the scalp was then sutured. The mice were placed in a heated cage and monitored until fully awake before being returned to their home cage for recovery.

### Behavior tests and memory engram labeling

Memory engram cell populations were labeled using the TRAP (targeted recombination in active populations) method. Following virus injection and recovery, cFos-CreER mice were administered tamoxifen (150 mg/kg) 24 hours prior to the behavioral testing and then returned to their housing cages. The mice underwent a contextual fear conditioning paradigm, which involved familiarizing them with a context (cubic shape, 75% alcohol scent) and subsequently exposing them to three trials of 0.5 mA foot-shocks (2 seconds per trial). After the conditioning, the mice were returned to their housing cages.

### Primary neuron culture

Cortical neuron cultures were prepared as previously described. Embryonic day 16 (E16) ICR mice were used for this process. The cortical regions of the embryos were meticulously dissected under a dissection microscope and then digested with 0.125% trypsin in the presence of 0.05% DNase I (Sigma). For imaging purposes, glass slides were coated with poly-D-lysine (PDL) and laminin, then washed with deionized water and air-dried. Following digestion, the tissue was dissociated into a single-cell suspension and seeded at a density of 10-15 × 10^5 cells per well in 12-well plates. The neurons were maintained in neural basal medium supplemented with 2% B27, 2 mM GluMax, and penicillin/streptomycin. Neurons were incubated at 37C and 5% CO2.

### Immunohistochemistry of mouse brain slices

Mice were anesthetized with 500 ul of 2% pentobarbital sodium and transcardially perfused with PBS, followed by 4% paraformaldehyde (PFA) dissolved in PBS. The brain tissue was carefully extracted from the skull and placed in a centrifuge tube containing PFA, then stored at 4°C overnight. To achieve optimal super-resolution imaging, tissue slicing and staining should be performed as soon as possible after fixation. Typically, brain sections are cut the following day into 10-15 µm slices, with the dorsal hippocampal region selected for subsequent processing. The brain slices were first incubated in 0.3% Triton X-100 for membrane permeabilization and then blocked in 5% bovine serum albumin (BSA) for 2-4 hours, with the blocking time adjusted based on staining results. Antibody staining was conducted in a 24-well plate with an antibody dilution buffer (1% Triton X-100 and 2% BSA), adding approximately 300 µL of antibody solution per well, with a maximum of two slices per well. Primary antibody incubation was performed at 4°C overnight, followed by secondary antibody incubation at room temperature for 1 hour. After antibody staining, the brain slices were transferred to a 12-well plate, with 2 mL of PBS per well for washing. The slices were washed three times, each for 15 minutes, to remove unbound antibodies and dyes and reduce background staining.

### Chemical LTP induction protocol

To artificially induce long-term potentiation (LTP), neurons cultured on coverslips were initially transferred to a Mg²⁺ free extracellular solution containing 150 mM NaCl, 3 mM KCl, 3 mM CaCl₂, 10 mM HEPES, 5 mM glucose, 0.5 μM tetrodotoxin, 1 μM strychnine, and 20 μM bicuculline methiodide, bicuculline methiodide purchased from MedChemExpress others are from sigma. The neurons were incubated in this solution in cell incubator for 10 minutes. Stimulation was applied by exposing the neurons to 100 μM glycine (Sigma) in the same Mg^2+^free solution for 5 minutes.

### Immunohistochemistry of cultured neuron sample

Neurons were fixed with 4% PFA for 10 minutes at room temperature, then washed three times with PBS, each for 5 minutes. Cells were permeabilized with 0.3% Triton X-100 for 5 minutes and blocked with 5% BSA for 25 minutes. Primary antibodies, prepared in 5% BSA, were added (60-70 µL per sample), and cells were incubated in a humidified chamber at 4°C overnight. The next day, cells were washed three times with PBS (5 minutes each), followed by secondary antibody staining in 5% BSA at room temperature for 1 hour. After staining, cells were washed three more times with PBS.

### Sample mounting and imaging buffers

To prepare all imaging samples in this study, approximately 120 µL of a PBS-diluted gold nanoparticle suspension was applied to the samples for incubation for 15 minutes following staining. After incubation, the samples were rinsed multiple times with PBS to remove any unbound nanoparticles. Next, 70-80 µL of imaging buffer was placed onto a glass slide with a single-sided groove, and the cover slip containing cultured neurons or brain slices was carefully placed over it, ensuring no air bubbles formed during placement. Excess buffer was wiped off, and the edges of the cover slip were sealed using a mixed silicone-glue solution. The imaging buffer used was a standard GLOX buffer, which consisted of 50 mM TRIS/HCl, 10 mM NaCl, 10% (w/v) glucose at pH 8.0, 64 µg/mL catalase, 0.4 mg/mL glucose oxidase, and 30mM 2-mercaptoethylamine (MEA).

### MINFLUX nanoscopy

MINLFUX imaging experiments were conducted using MINFLUX nanoscope from Abberior Instruments, equipped with a 100 × Oil immersion objective lens (NA 1.4). The system integrates a 642-nanometer continuous-wave excitation laser and a 405-nanometer continuous-wave activation laser, alongside a spatial light modulator for precise beam shaping. The imaging setup includes an electro-optical detector-based MINFLUX scanner and two avalanche photodiodes, coupled with fluorescence filters (650 to 750 nanometers) to detect emitted fluorescence.To identify region of interest, we utilized the equipped confocal module with a 642-nanometer laser. ROIs were selected based on these confocal images. For subsequent MINFLUX imaging, we activated the MINFLUX642 laser channel and chose 3D MINFLUX modes for data acquisition.The imaging system was stabilized using a dedicated stabilization module to minimize fluctuations. Nanoscale gold particles were incorporated as fiducial markers to enable continuous monitoring and correction of any drift or instability in the imaging system. This stabilization ensured that the MINFLUX system maintained consistent XYZ axis precision, with fluctuations restricted to within 1 nanometer throughout the imaging process, thereby ensuring the reliability and accuracy of the imaging data.

### MINFLUX localizations data processing and clustering

Localization data from the MINFLUX imaging were processed and analyzed using custom MATLAB scripts. The data processing workflow involved loading raw data, noise removal, clustering, and visualization, as outlined below. Data Loading and Preprocessing: Localization data were loaded from.mat files containing the coordinates (X, Y, Z) of the MINFLUX-localized points. The area number was extracted from the file name, and the corresponding data were loaded for further analysis. The raw data were then scaled by a factor of 3 to convert the units to nanometers. Clustering Using DBSCAN: DBSCAN (Density-Based Spatial Clustering of Applications with Noise) was employed to group localized points into spatial clusters. The algorithm was configured with an epsilon (ε) parameter to define the neighborhood radius and a minPoints parameter to set the minimum number of points required to form a cluster. Points classified as noise (with an index of-1) were removed from further analysis. Data Visualization: To visualize the data before and after clustering, 3D scatter plots were generated. Initially, the raw localizations were plotted to show their spatial distribution. After DBSCAN clustering, the points were color-coded according to their assigned cluster index to visualize the clustering results.

### Local Density Calculation

To calculate the local density for each cluster, the centroids of the clusters were first determined. The centroid of each cluster was computed as the mean of all points within the cluster. For each cluster, the points belonging to it were identified based on their cluster index, and the mean coordinates of these points were calculated and scaled to nanometer units.Following centroid determination, a local density map was generated by calculating the number of other centroids within a specific radius, R. This radius was set to 150 nm. For each centroid, the distances to all other centroids were computed, and the number of centroids within the specified radius was counted. The local density for each centroid was then normalized by dividing the count of nearby centroids by the volume of a sphere with radius R, which was calculated as 4πR^3^/3. The resulting normalized local density values were used to visualize the density distribution of the cluster centroids in 3D, with color-coding based on the density values.

## Acknowledgement

The work was supported by the Shanghai Frontiers Science Center Program (2021–2025 No. 20) to M.G., the Science and Technology Commission of Shanghai Municipality (Grant No. 21DZ1100500), the Shanghai Municipal Science and Technology Major Project to M.G., the Science & Technology Innovation 2030 Project of China (2021ZD0203501) to J.-.S.G., and Natural Science Foundation of China(32130043) to H.X..

## Contributions

M.G., H.X., and J-S.G. conceived and designed the project. Y.H. performed the experimental data collection with help from X.W. and S.X. Y. H led the data analysis with assistance from K. L. The manuscript was written by Y. Hu and and all authors contributed to the final manuscript.

## Supplementary materials

**Supplementary figures (S1-S8)**

**Supplementary tables(S1-S2)**

---

**Supplementary Fig. 1.**
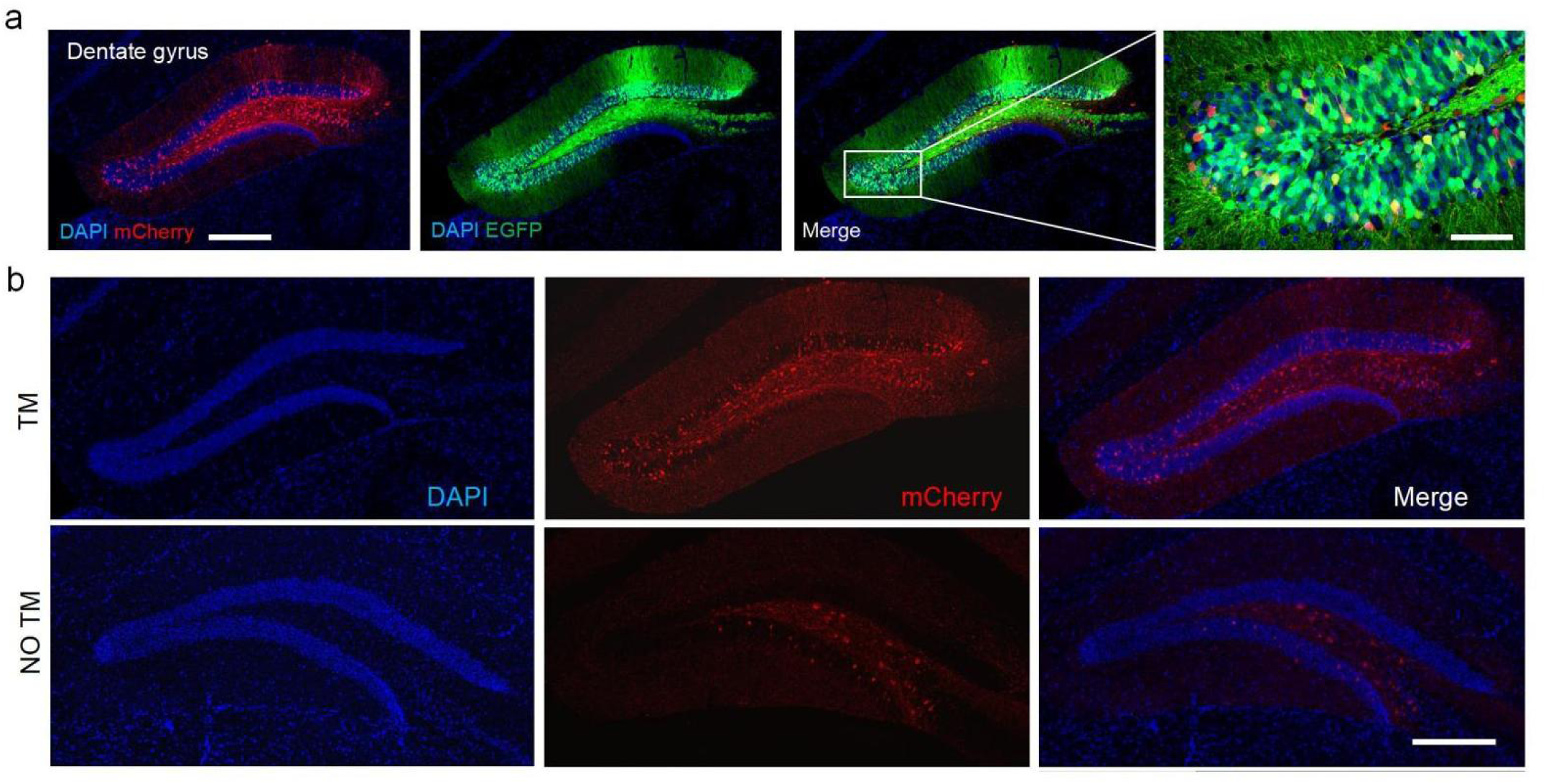
Activity-dependent labeling of neurons in hippocampal DG region. **a.** Complete confocal image of hippocampal DG region immunofluorescence labeling. The blue channel represents the nuclear stain DAPI, red represents mcherry, and green represents EGFP. Scale bar: 500 μm. **b.** Activity-dependent labeling in the hippocampal region in the presence or absence of TM injection. The blue channel represents DAPI (nuclear stain), and red represents mcherry.Scale bar: 500 μm.

**Supplementary Fig. 2.**
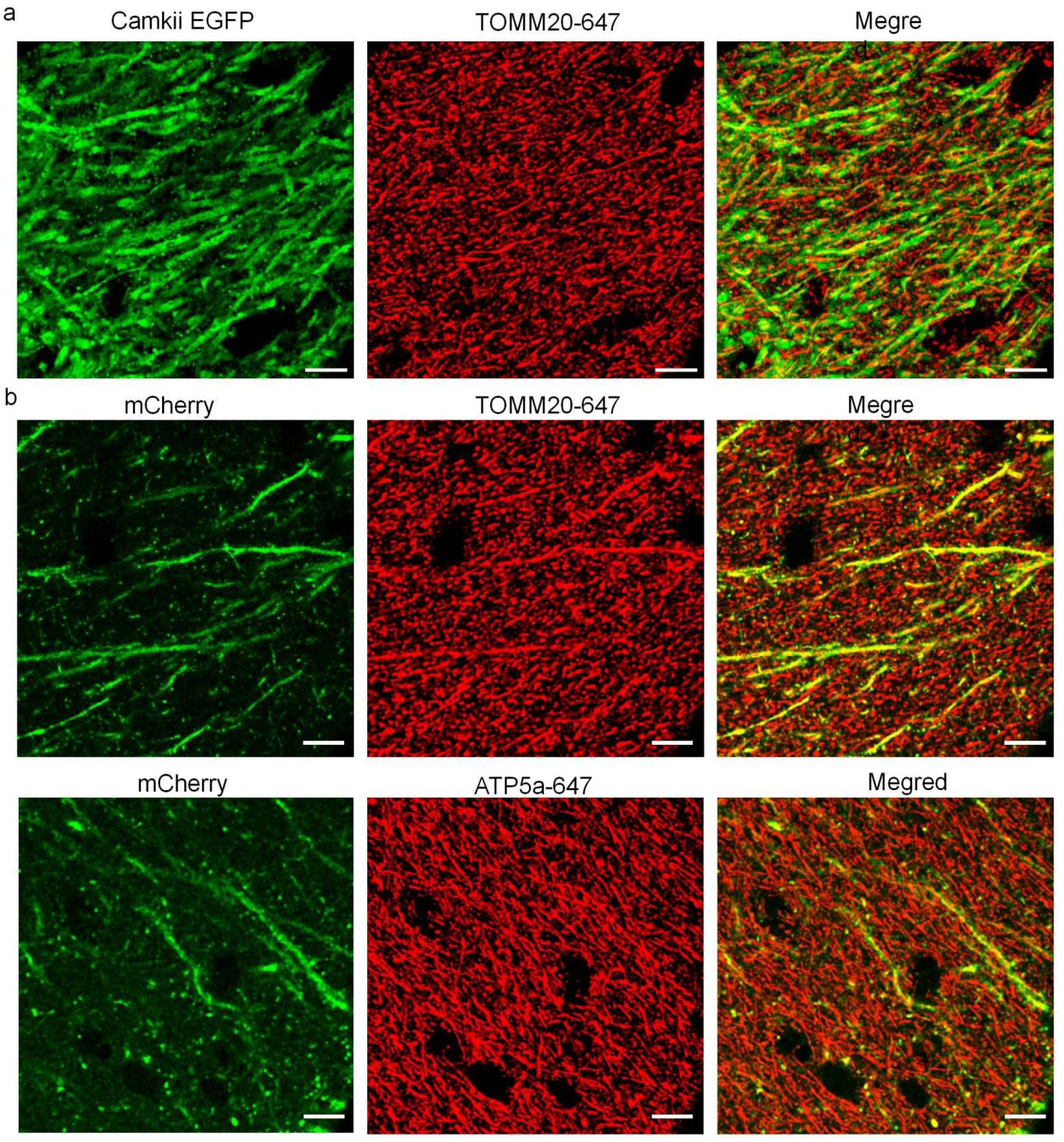
Mitochondrial morphology in DG molecular layers. **a.** Mitochondria in DG Camkii-positive cells in the molecular layer dendrites. Mitochondria were labeled with TOMM20 primary antibody conjugated to AF647 dye, showing red (TOMM20-647), and EGFP was used to label the cells, showing green.**b.** Mitochondria in DG memory engram cells in the molecular layer dendrites. Mitochondria were labeled with TOMM20 primary antibody conjugated to AF647 dye and ATP5a primary antibody conjugated to AF647 dye respectively. Scale bar: 10 μm.

**Supplementary Fig. 3.**
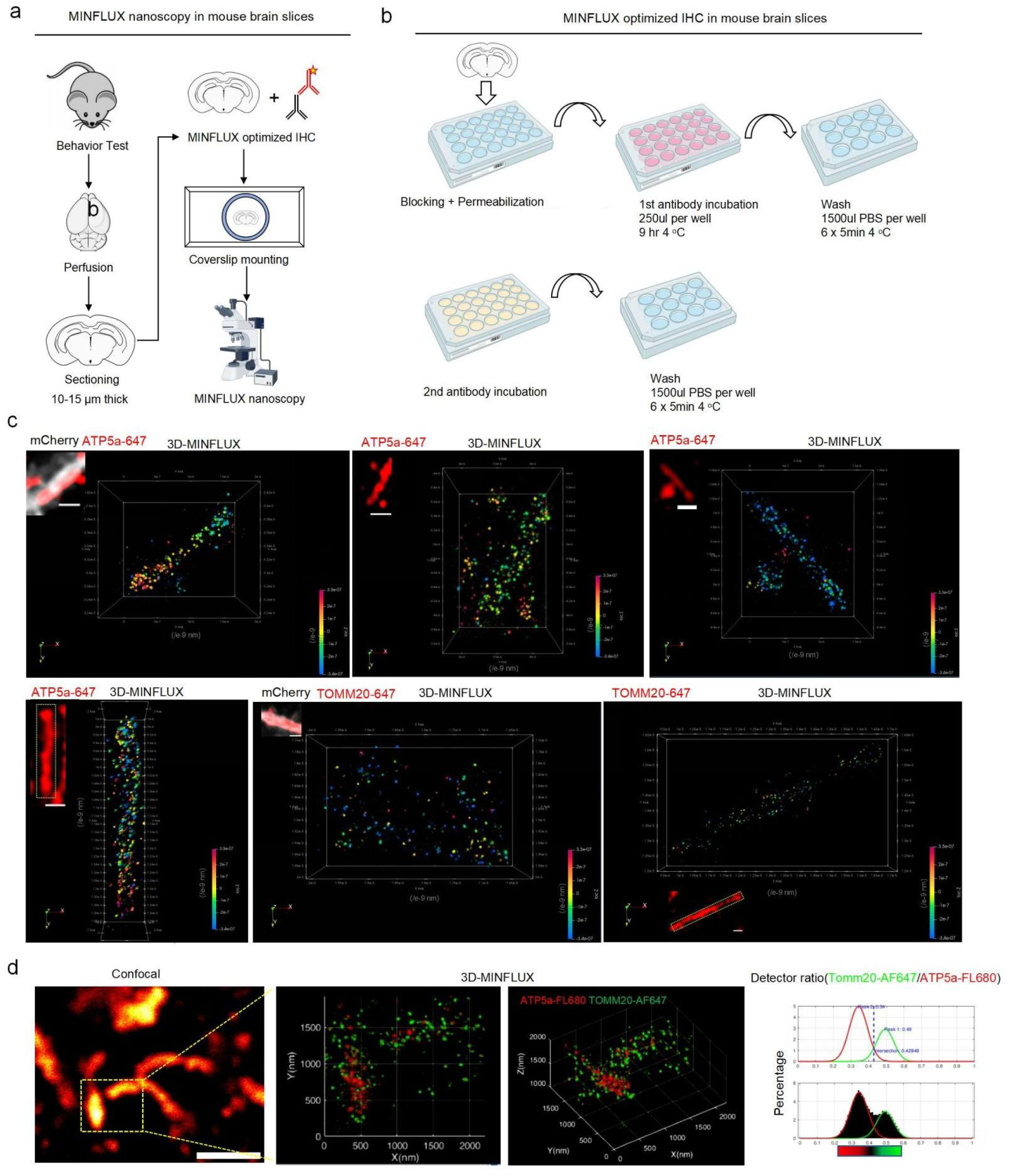
Brain section MINFLUX staining protocol and 3D rendering MINFLUX imaging results. **a.** MINFLUX brain slice super-resolution imaging workflow (see Methods section for details). **b.** Optimized brain slice MINFLUX imaging protocol, including membrane permeabilization, blocking, and extended washing with PBS. **c.** Three-dimensional MINFLUX imaging of TOMM20 and ATP5a proteins in mitochondria, using primary antibodies directly conjugated to AF647 dye. Individual localizations were visualized as 3D Gaussian functions with a σ value of 3 nm. Data were processed and visualized using ParaView software. **d.** Dual-color 2D and 3D MINFLUX imaging of TOMM20 and ATP5a proteins in brain slices. ATP5a protein was labeled with a primary antibody and then stained with a secondary antibody conjugated to FL680. TOMM20 was stained using a primary antibody conjugated to AF647.Scale bar: 1 μm.

**Supplementary Fig. 4.**
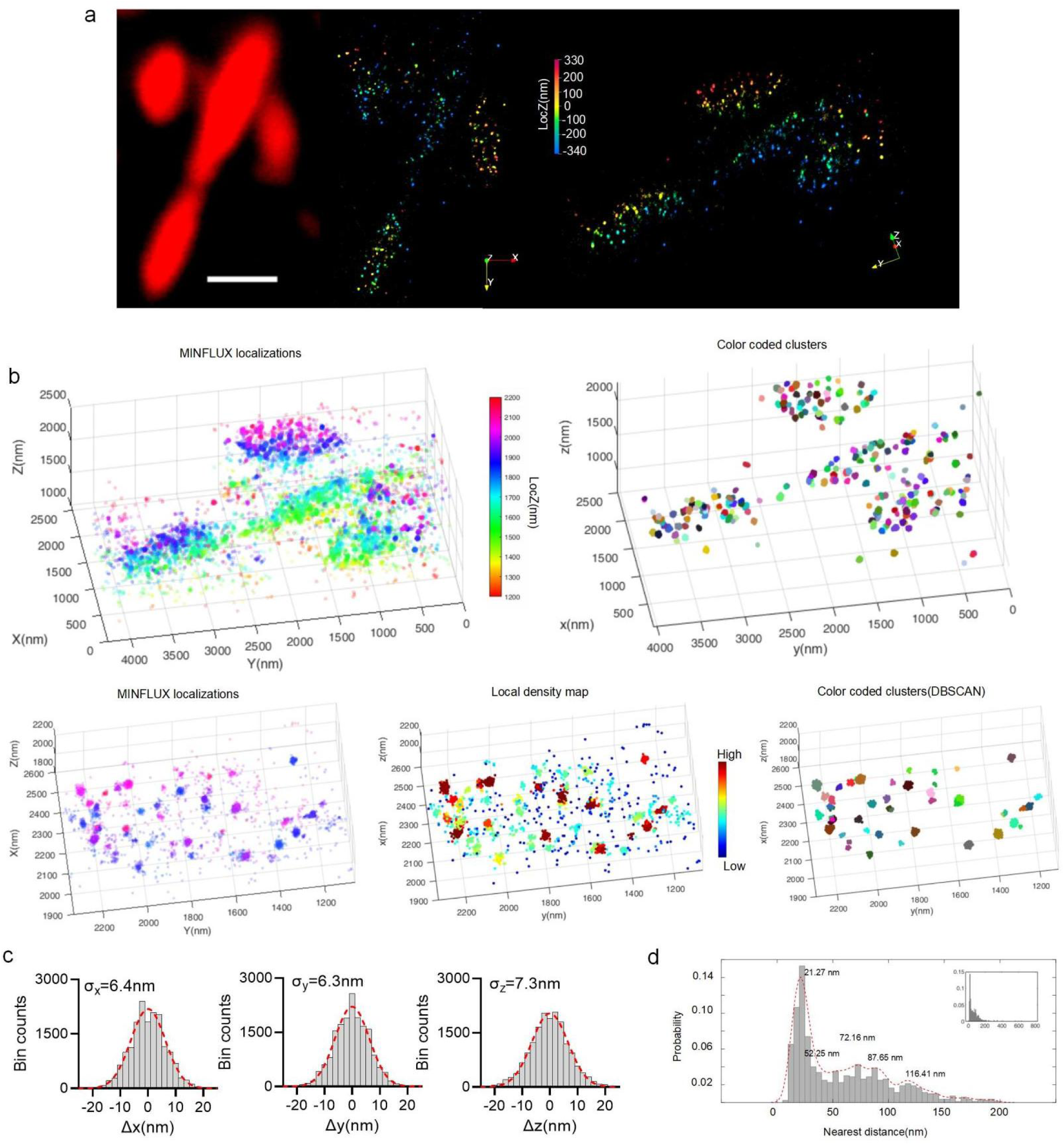
Molecular assignment based on MINFLUX localization data. **a.** 3D MINFLUX imaging of ATP5a protein in mitochondria in a brain section experiment, Scale bar: 500 nm.. **b.** Upper panel: 3D MINFLUX imaging data of ATP5a protein in a brain slice (left) and clustering results based on density-based segmentation (right), with each cluster represented in a different color. Lower panel: Local 3D MINFLUX data showing the localization density map. The hot spots of each cluster are marked, and clustering was performed using the DBSCAN algorithm in MATLAB with minpoints set to 12 and epsilon set to 8. **c** Statistical analysis of localization precision along the x, y, and z axes. The distribution data were fitted using Gaussian fitting, and the standard deviation (σ) was calculated. The result of Gaussian fitting is indicated by the red dashed line.**d** The nearest distances between centroids were calculated and the distribution is presented. The inset shows the distribution of all nearest distances, with the main figure representing data within the 0–200 nm range.

**Supplementary Fig. 5.**
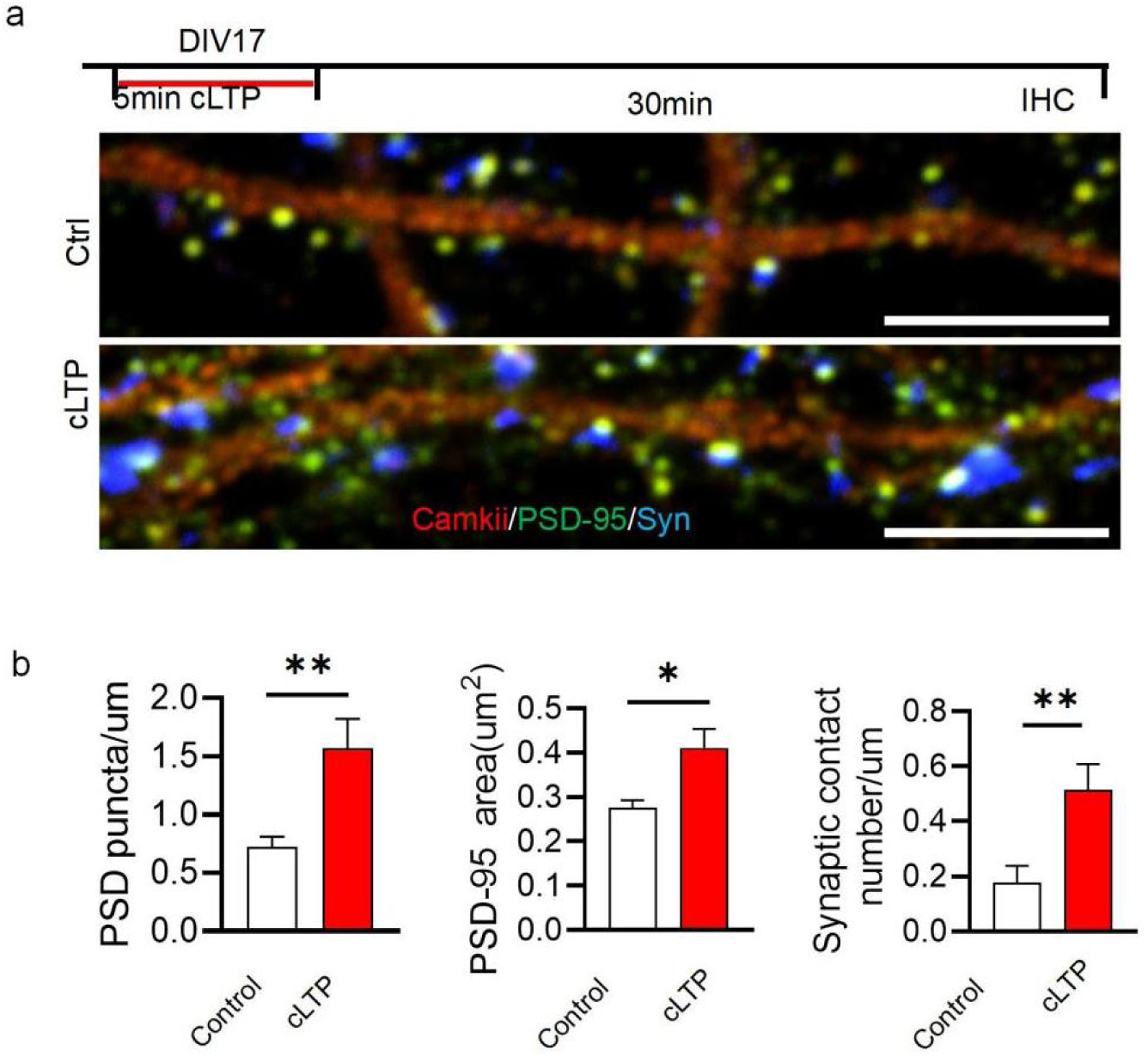
cLTP-induced PSD95 signal enhancement in synapses. **a.** Induction of cLTP in DIV17 neuronal cultures. Immunofluorescence staining for PSD95 (green) and camkii (red) was performed 30 minutes after induction, with the blue channel showing virus-induced syn expression. **b.** Analysis of PSD puncta density, size, and the density of PSD95-syn pairing. Unpaired t-test, *P < 0.05, **P < 0.01.

**Supplementary Fig. 6.**
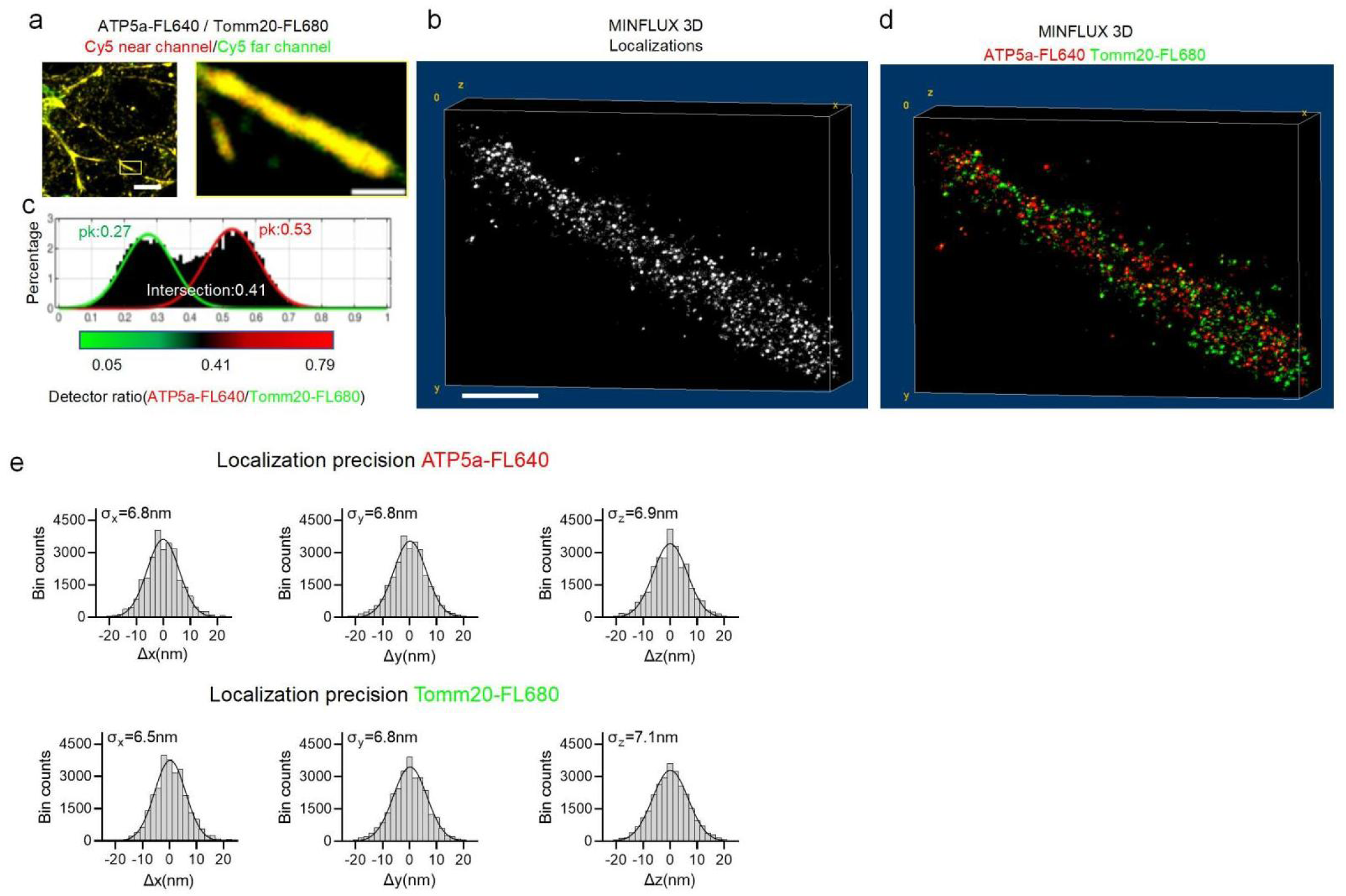
Localization precision of mitochondrial dual-color MINFLUX imaging. **a,d** Dual-color MINFLUX imaging of mitochondrial inner and outer membrane proteins in neurons.**a** Mitochondrial inner membrane protein ATP5a and outer membrane protein TOMM20 were labeled with primary antibodies and subsequently visualized using secondary antibodies conjugated to FL640 and FL680, respectively. ATP5a signals were acquired using the Cy5 near channel, while TOMM20 signals were collected through the Cy5 far channel and pseudo-colored red and yellow, respectively. Scale bar: 10 um (large image), 1 um (small image).**b** Localization data from dual-color MINFLUX imaging, with 3D volumes displayed using ImageJ’s Volume Viewer.**c** Distribution of detector ratios for each localization. The critical points for FL640 and FL680 assignments were determined using double Gaussian fitting, with the intersection of the two Gaussian curves marking the threshold for distinguishing the fluorophores. **d** Dual-color 3D MINFLUX imaging of inner membrane protein ATP5a and outer membrane protein TOMM20, after separating FL640 and FL680 signals. Scale bar: 400 nm. **e.** Localization precision data along the x, y, and z axes for FL640 and FL680 fluorophores during 3D MINFLUX imaging. The black curves represent Gaussian fits, and SD values were calculated from the fitting results.

**Supplementary Fig. 7.**
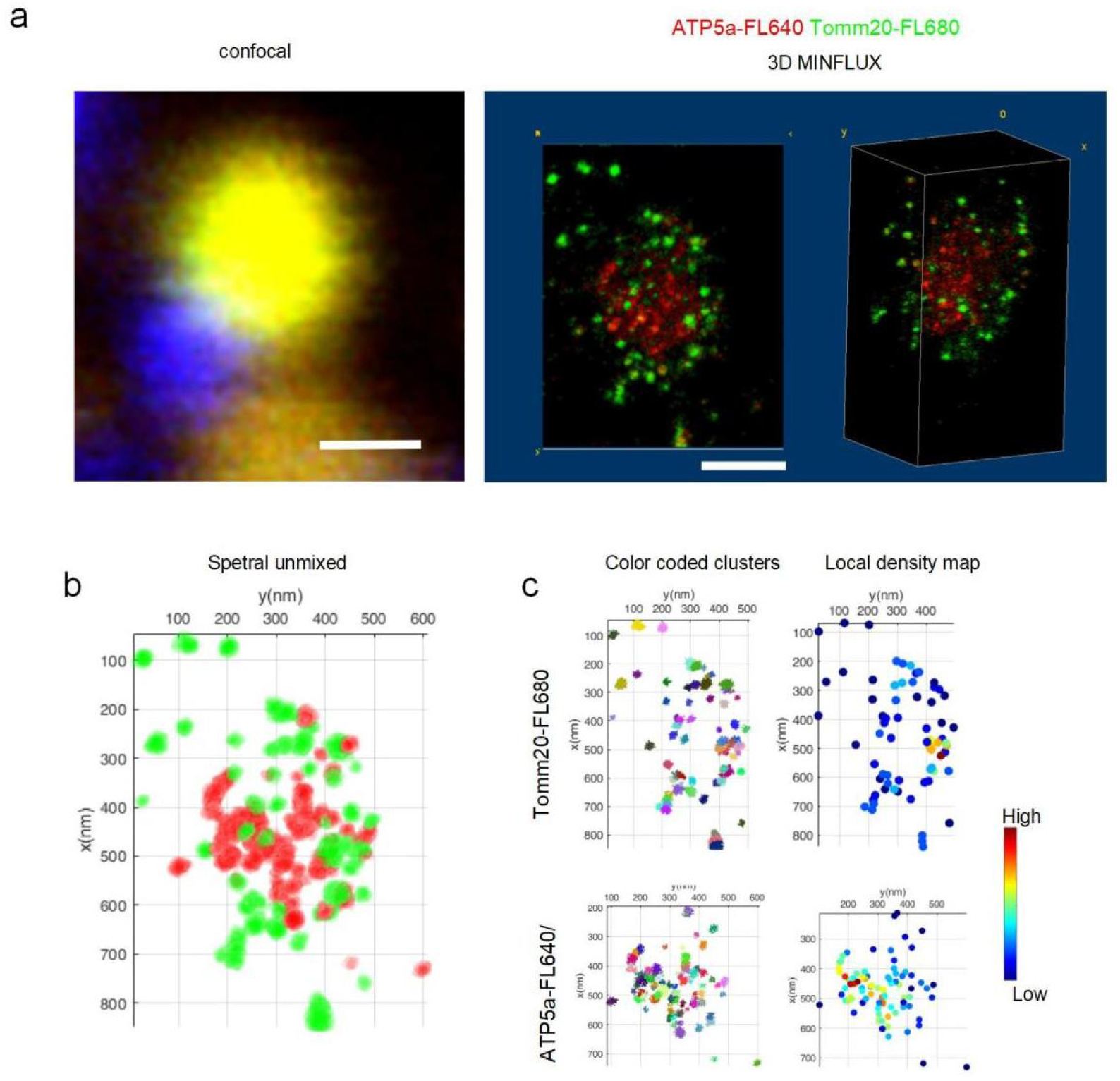
Local density processing for dual color 3D MINLUX imaging. a. A 3D MINFLUX imaging result of mitochondria near pre-synaptic syn signals. In the confocal image, red indicates ATP5a, and green indicates TOMM20, both labeled with primary antibodies and then stained with secondary antibodies conjugated to FL640 and FL680, respectively. The 3D MINFLUX data is presented with the same color scheme: red for ATP5a and green for TOMM20. The left side shows the z-axis projection, and the right side shows the 3D view, both generated using the Volume Viewer plugin in ImageJ. Scale bars: confocal, 500 nm; MINFLUX, 200 nm. b. Digitally displayed separation of the two dyes based on detector ratio in MATLAB, followed by DBSCAN clustering of the localizations for both dyes. The local density for each cluster was calculated based on the centroids for further analysis. c. Example of the DBSCAN clustering results and local density calculations for the TOMM20-FL680 and ATP5a-FL640 channels.

